# *miR-206* family is important for mitochondrial and muscle function, but not essential for myogenesis *in vitro*

**DOI:** 10.1101/796821

**Authors:** Roza K. Przanowska, Ewelina Sobierajska, Zhangli Su, Kate Jensen, Piotr Przanowski, Sarbajeet Nagdas, Jennifer A. Kashatus, David F. Kashatus, Sanchita Bhatnagar, John R. Lukens, Anindya Dutta

**Affiliations:** Department of Biochemistry and Molecular Genetics, University of Virginia School of Medicine, Pinn Hall 1232, Charlottesville, Virginia 22908, USA; Department of Microbiology, Immunology and Cancer Biology, University of Virginia School of Medicine, Charlottesville, Virginia 22908, USA; Department of Neuroscience, University of Virginia School of Medicine, Charlottesville, Virginia 22908, USA; Center for Brain Immunology and Glia, Department of Neuroscience, School of Medicine, University of Virginia, Charlottesville, Virginia 22908, USA

**Keywords:** myomiRs, skeletal muscle differentiation, myogenesis, miR-1a-1, miR-1a-2, miR-206, embryonic lethality, mitochondria function, muscle function

## Abstract

*miR-206*, *miR-1a-1* and *miR-1a-2* are induced during differentiation of skeletal myoblasts and promote myogenesis *in vitro*. *miR-206* is required for skeletal muscle regeneration *in vivo*. Although this microRNA family is hypothesized to play an essential role in differentiation, a triple knockout of the three genes has not been done to test this hypothesis. We report that triple KO C2C12 myoblasts generated using CRISPR/Cas9 method differentiate despite the expected de-repression of the microRNA targets. Surprisingly, their mitochondrial function is diminished. Triple KO mice demonstrate partial embryonic lethality, most likely due to the role of miR-1a in cardiac muscle differentiation. Two triple KO mice survive and grow normally to adulthood with smaller myofiber diameter and diminished physical performance. Thus, unlike other microRNAs important in other differentiation pathways, the *miR-206* family is not absolutely essential for myogenesis and is instead a modulator of optimal differentiation of skeletal myoblasts.

## Introduction

The downregulation of pluripotency markers and activation of lineage-specific gene expression during differentiation allow for accurate development. Differentiation-induced microRNAs play a major role in this process by repressing their targets – genes responsible for self-renewal. Depletion of DGCR8 protein essential for biogenesis of microRNA in pluripotent cells decreases most active microRNA levels and inhibits differentiation (Wang et al., 2007). Since many microRNAs are induced during differentiation of specific tissue lineages, several of them have been tested for their importance in differentiation, particularly whether they act as a switch that is essential for differentiation, or as a modulator of differentiation. MicroRNAs regulate processes as early as gastrulation (Choi et al., 2007; Rosa et al., 2009), neural development (Delaloy et al., 2010; Krichevsky et al., 2006; Zhao et al., 2009), muscle development (Chen et al., 2006; Cordes et al., 2009; Dey et al., 2012; Sarkar et al., 2010; Zhao et al., 2005), bone formation (Li et al., 2009; Li et al., 2008), skin development (Jackson et al., 2013; Wang et al., 2013) and hematopoiesis (Chen et al., 2004; Garzon and Croce, 2008; Zhu et al., 2013). Many of the studied microRNAs have been suggested to be essential, e.g. *miR-206* and *miR-1a* for skeletal muscle myoblast differentiation (Chen et al., 2010; Dey et al., 2011), *miR-144/451* for erythroid cells differentiation (Dore et al., 2008; Rasmussen et al., 2010), *miR-17∼92* during B lymphopoiesis and lung development (Ventura et al., 2008), *miR-15a-1* and *miR-18a* for development and function of inner ear hair cells in vertebrates (Friedman et al., 2009), *miR-219* for normal oligodendrocyte differentiation and myelination (Dugas et al., 2010), *miR-204* for differentiation of the retinal pigmented epithelium (Ohana et al., 2015) and *miR-375* for human spinal motor neuron development (Bhinge et al., 2016).

Myogenesis is a process of muscular tissue formation, which first occurs in vertebrate embryonic development (Parker et al., 2003), but also happens in adult muscle regeneration (Chargé and Rudnicki, 2004). The skeletal muscle satellite cells are the myogenic stem cells of adult muscles residing between the sarcolemma and basal lamina of muscle fibers (Mauro, 1961). In normal conditions these tissue specific stem cells stay in a quiescent G0 state (Cheung and Rando, 2013). Upon activation by injury or disease, they re-enter the cell cycle to establish a population of skeletal muscle progenitors (myoblasts), which differentiate further and fuse to produce myotubes. The major regulator of this process is *Pax7* transcription factor (Olguín and Pisconti, 2012; Zammit et al., 2006). The satellite cells express the transcription factor Pax7 in G0 and when activated, coexpress *Myod1*. Downregulation of *Pax7* leads to differentiation into myotubes, whereas downregulation of *Myod1* leads to a return to quiescence. An important player in *Pax7* downregulation and differentiation induction is *miR-206* (Chen et al., 2010; Dey et al., 2011). Other microRNAs are also known to play important roles in skeletal muscle differentiation. A conditional skeletal muscle specific knockout of Dicer in mice leads to global loss of miRNAs in developing skeletal muscle, resulting in widespread apoptosis and abnormal myofiber morphology (O’Rourke et al., 2007). Over the last 14 years *miR-206*, *miR-1a-1* and *miR-1a-2* have been hypothesized to not only be very important for muscle differentiation, but also essential for this process (Anderson et al., 2006; Chen et al., 2006; Chen et al., 2010; Dey et al., 2011; Gagan et al., 2012; Goljanek-Whysall et al., 2012; Heidersbach et al., 2013; Hirai et al., 2010; Kim et al., 2006; Koutsoulidou et al., 2011; Kwon et al., 2005; Mishima et al., 2009; Rao et al., 2006; Sokol and Ambros, 2005; Sweetman et al., 2008; Vergara et al., 2018; Wystub et al., 2013; Wüst et al., 2018; Yuasa et al., 2008; Zhao et al., 2007; Zhao et al., 2005). *miR-206*, *miR-1a-1* and *miR-1a-2* are members of the myomir family and are expressed from bicistronic loci. Interestingly, *miR-206* and *-1a* have an 18/21 base match in sequence with each other and complete identity in the first eight nucleotides that constitute the seed sequence for target recognition. Even though all three are expressed in skeletal muscles, *miR-1a-1* and *miR-1a-2* are also expressed in cardiac muscle, where *miR-206* is not expressed (Kim et al., 2006; Sempere et al., 2004). All three are upregulated during murine skeletal myoblast differentiation (Kim et al., 2006). Overexpression of *miR-206* induces C2C12 differentiation, whereas simultaneous knockdown of *miR-206* and *miR-1a* results in diminished differentiation (Kim et al., 2006). Based on this it was hypothesized that the three microRNAs collectively are essential for skeletal muscle differentiation.

Knockout of the *miR-206* gene produced viable mice with no defect in skeletal muscle development, although there was a defect in skeletal muscle regeneration after extensive muscle injury (Liu et al., 2012). The loss of *miR-206* in mice modeling amyotrophic lateral sclerosis leads to faster disease progression and lack of regeneration of neuromuscular synapses, again suggesting a role of the microRNA in response to tissue injury. In contrast, there was high embryonic lethality of *miR-1a-2* knockout mice caused by various cardiac problems (Zhao et al., 2007). Although *miR-1a-1* knockout showed embryonic lethality in the 129 genetic background animals, knockout pups were obtained at the expected frequency in a mixed genetic background, proving that the *miR-1a-1* knockouts can produce viable mice (Heidersbach et al., 2013). Although the double KO of *miR-1a-1* and *miR-1a-2* was found to have high embryonic lethality, some mice survived with normal skeletal muscle development. By 10th postnatal day, however, all double knockout animals were dead because of serious cardiac defects (Heidersbach et al., 2013). Moreover, knockout of two bicistronic loci (*miR-1a-1/133a-2* and *miR-1a-2/133a-1 dKO*) showed complete embryonic lethality at embryonic stage E11.5 (Wystub et al., 2013). The *miR-1a/133a* skeletal muscle specific dKO mice were alive, but had metabolic problems caused by incorrect functioning of mitochondria (Wüst et al., 2018).

Overall, all animals had relatively normal skeletal muscle development, and this could be explained because at least one myomir remained intact in all these animals. A complete knockout of all three microRNAs, *miR-206*, *miR-1a-1* and *miR-1a-2*, tests whether an essential role of these myomiRs in myogenesis was masked by the redundant function of the microRNA left in the cells. Here, we show that *miR-206*, *miR-1a-1* and *miR-1a-2* are not essential for skeletal muscle differentiation, but they are required for normal muscle formation and function. By analyzing triple KO C2C12 clones, we demonstrate that the myoblasts differentiate, although with decreased mitochondrial function. Additionally, triple KO animals reveal the three microRNA are not essential for muscle formation, but their absence causes defects in physical performance and myofiber width. Our results indicate that although *miR-206*, *miR-1a-1* and *miR-1a-2* are important for skeletal muscles and especially for their optimal function, they are not essential for myogenesis. Thus, this microRNA family, first discovered because of their induction during differentiation of stem cells and their ability to induce differentiation-specific genes, is not essential to trigger a differentiation switch, but is involved in modulating differentiation. In addition, it is surprising that the *miR-206* family and *miR-133a* are independently required for mitochondrial function even though they repress different gene targets.

## Results

### miR-206, miR-1a-1 and miR-1a-2 are not essential for C2C12 myoblasts differentiation

To determine whether *miR-206* and *miR-1a* are necessary for murine myoblast proliferation and differentiation, we designed 6 different sgRNAs to delete all three genes at once (Figure 1A, Table S1). To increase knockout efficiency, we used C2C12 cell line with stable overexpression of Tet-inducible Cas9. miRNA gene deletions were confirmed by PCR on genomic DNA (Figure 1B) and Sanger sequencing of PCR products (Figure 1C). We didn’t detect any pre-miRNA or miRNA left in our triple KO (tKO) clones (Figure 1D and S1A). To ensure the reproducibility of our results we obtained triple KO clones independently from PAX7 negative (Figure 1A-D) and PAX7 positive (Figure S1) C2C12 myoblasts.

**Figure 1.**
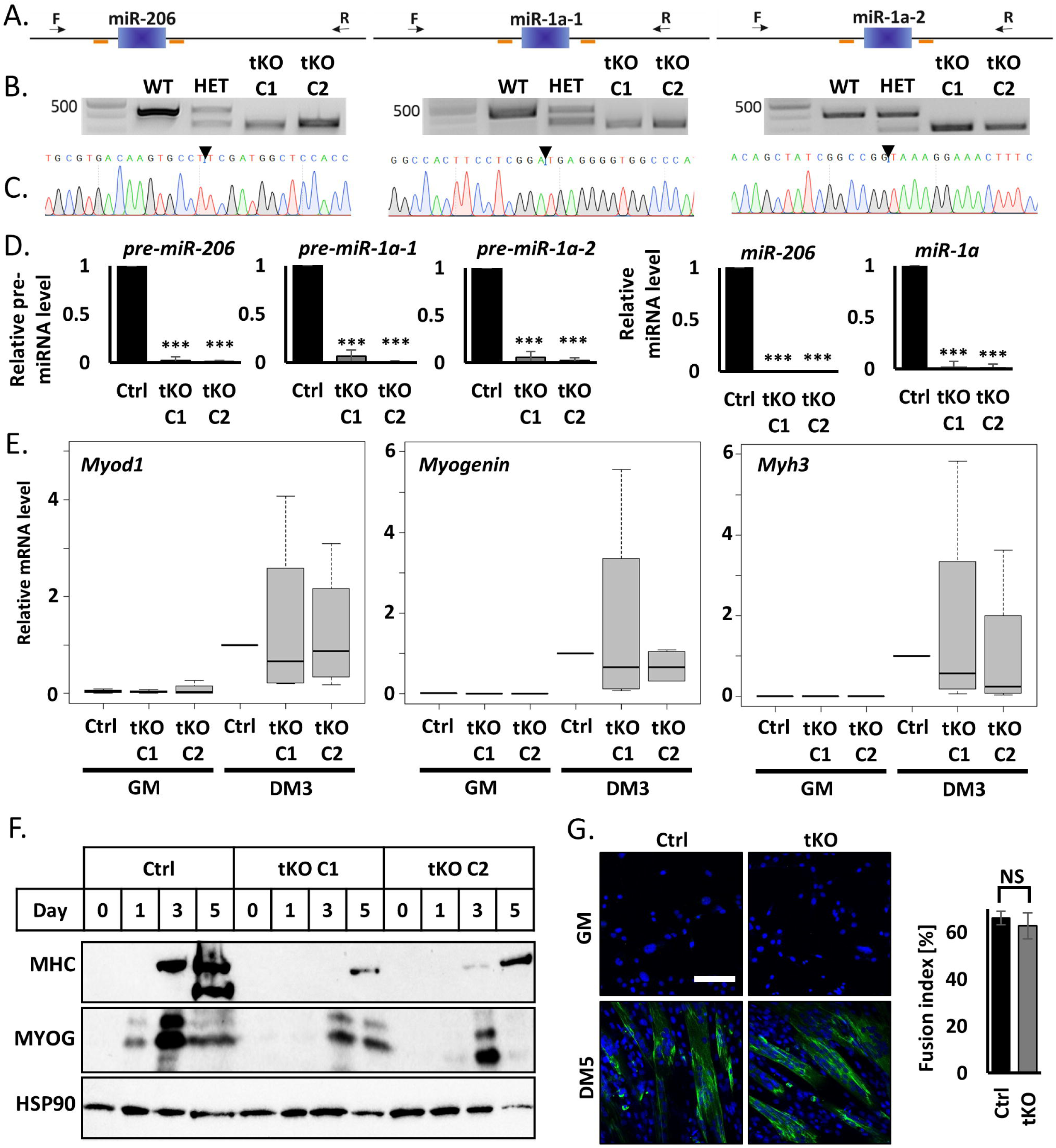
Simultaneous knockout of *miR-206*, *miR-1a-1* and *miR-1a-2* in C2C12 murine myoblast does not block cell differentiation. A) Scheme of CRISPR/Cas9 experiment design. Blue blocks represent genes, orange lines sgRNAs sequences and black arrow genotyping primers localization. Left: *miR-206*, middle: *miR-1a-1*, right: *miR-1a-2*. B) Representative picture of PCR genotyping results of triple KO C2C12 cells. Left: *miR-206*, middle: *miR-1a-1*, right: *miR-1a-2*. Top band – wild-type size, bottom band – KO size. C) Representative picture of Sanger sequencing confirmation of the genotyping PCR product in tKO C1 C2C12 clone. D) qRT-PCR analysis of differentiating (DM3) control cells (Ctrl) and triple KO clones (tKO C1 and tKO C2). Levels of pre-miRNAs were normalized to *Gapdh* and miRNAs – to *U6* snRNA. Levels are shown relative to control cells (Ctrl DM3). Values represent three biological replicates, presented as mean +/− SEM. Statistical significance was calculated using t-student test. (***) indicates p-value =< 0.001. E) qRT-PCR analysis of proliferating (GM) and differentiating (DM3) control cells (Ctrl) and triple KO clones (tKO C1 and tKO C2). Levels of *Myod1*, *Myogenin* and *Myh3* mRNAs were normalized to *Gapdh* and shown relative to control cells (Ctrl DM3) in box and whiskers plots. Values represent four biological replicates, black line represents median. Statistical significance was calculated using t-student test. F) Representative Western blot of time course of differentiation (GM, DM1, DM3, DM5) of control cells (Ctrl) and triple KO clones (tKO C1 and tKO C2). Protein levels for MYOGENIN and MHC were measured. HSP90 serves as a loading control. G) Immunofluorescence analysis of fixed cells 5 days after differentiation (DM5). Cells were immunostained with antibodies against MHC. Hoechst 33342 was used to visualize nuclei. Quantification of the fusion index is presented on the right side. White line = 200µm.

C2C12 clones lacking *miR-206* and *miR-1a* differentiate upon serum starvation producing similar levels of promyogenic mRNAs: *Myod1, Myogenin* and *Myh3* as control cells (Figure 1E). The tKO clones also produce the corresponding proteins, which include markers of early (MYOGENIN) and late myogenesis (MHC), though their levels are lower in the PAX7 negative tKO cells (Figure 1F). In PAX7 positive tKO clones the *Myod1*, *Myogenin* and *Myh3* mRNAs are induced normally during differentiation (Figure S1B), but we observe similar levels of MYOGENIN protein and slightly decreased MHC protein (Figure S1C). Moreover, the fusion index counted in the PAX7 positive cells (Figure 1G) shows there is no morphological differences upon differentiation between the control and tKO cells.

### Although triple KO C2C12 clones differentiate they are impaired in metabolic performance

By RNA-seq the triple KO clones differentiated efficiently as confirmed by hierarchical clustering which shows the differentiating tKO cells cluster with the differentiating wild type cells, separate from proliferating cells (Figure 2A). Nevertheless, 531 and 412 genes were differentially expressed in tKO in comparison to control cells in proliferating and differentiating conditions, respectively (Figure 2B). Among 232 genes upregulated during differentiation in the tKO cells are several involved in retinoic acid regulation (from the top 10: *Aldh1a1*, *Aldh1a7*, *Brinp3* and *Tceal5*), which was previously described as important factor for myogenesis (El Haddad et al., 2017; Halevy and Lerman, 1993; Lamarche et al., 2015). Aldehyde dehydrogenases (*Aldh*) are also known to have promyogenic potential (Jean et al., 2011; Vauchez et al., 2009; Vella et al., 2011).

**Figure 2.**
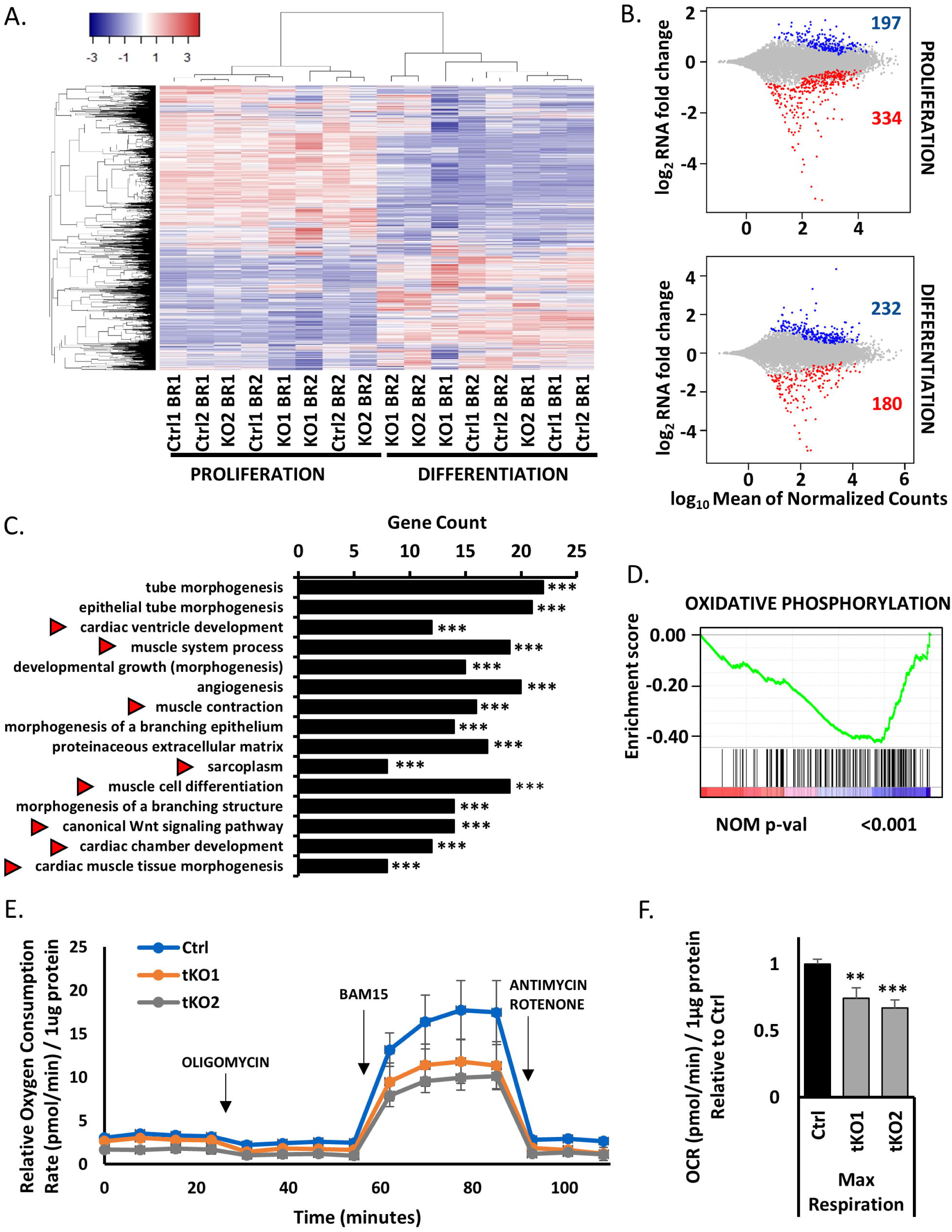
RNAseq analysis confirms the tKO cells differentiate, but they have lower mitochondrial respiratory capacity than control cells during differentiation. A) Heatmap showing clustering of all RNA-seq samples and conditions based on expression (FPKM≥1) in proliferating conditions (left) and in differentiating conditions (right) in control cells (Ctrl1 and Ctrl 2) and triple KO clones (KO1 and KO2). There are two biological replicates for each condition (BR1 and BR2). B) MA plots representing differentially expressed genes between control cells and triple KO clones in proliferation (top) and differentiation conditions (bottom). Upregulated genes are presented in blue and downregulated in red (p_adj_<0.01). C) Top 15 significant Gene Ontology terms enriched in downregulated genes in differentiating triple KO clones (tKO) in comparison to differentiating control cells (Ctrl). Red arrows show gene terms related to muscle development and regeneration. (***) indicates p_adj_ =< 0.001 D) Enrichment plot from gene set enrichment analysis (GSEA) showing the gene set involved in oxidative phosphorylation is enriched among differentially downregulated genes in differentiating tKO clones versus differentiating control cells. E) Oxygen Consumption Rate (OCR) of differentiating (DM1) tKO C2C12 clones and control C2C12 cells measured at basal conditions and after the administration of the indicated compounds. Values represent five biological replicates, presented as mean +/− SEM. Levels are shown relative to the total protein content. F) Maximal Respiration of control C2C12 cells and tKO clones in differentiation (DM1). Values represent five biological replicates, presented relative to control cells as mean +/− SEM. (**), (***) indicates p_adj_ < 0.01 and 0.001 respectively. Levels are shown relative to the total protein content and control cells.

Gene Ontology (GO) analysis of differentiating triple KO clones reveals downregulation of genes involved in skeletal and cardiac muscle development pathways (Figure 2C). GO also shows that genes involved in mitochondria biogenesis and function are induced during differentiation of WT control cells, but not in the tKO cells (Figure S2A, B). The latter is supported by Gene Set Enrichment Analysis (GSEA), which shows only one significant category – an enrichment of genes related to oxidative phosphorylation among the genes downregulated in differentiated tKO cells vs. differentiated control cells (Figure 2D).

To assess whether tKO perturbed the mitochondrial metabolism of C2C12 cells, we analyzed oxygen consumption rate (OCR) as a measure of aerobic respiration using the Seahorse Bioscience XF24. The tKO clones had a lower OCR even at baseline than the control cells in differentiation conditions (Figure 2E). Furthermore, treatment of these clones with the mitochondrial uncoupling agent BAM15, which produces maximal oxygen consumption, revealed an even greater difference in OCR between control and tKO clones (Figure 2F). These results suggest that these three microRNA are required to maintain mitochondrial respiratory capacity, and are consistent with the findings from *miR-1a-/-133a-/-* skeletal muscle specific double KO animals that showed impairment of mitochondrial function (Wüst et al., 2018). Thus, although the tKO C2C12 clones differentiate, their differentiation is slightly impaired compared to the WT cells and the cells suffer a downregulation of mitochondria function.

### miR-206 and miR-1a targets are specifically upregulated during differentiation of triple KO C2C12 cells

The absence of *miR-206* and *miR-1a* could be compensated by some other mechanism, in which case the targets of *miR-206* and *miR-1a* will be unchanged in the tKO cells. We therefore analyzed the expression levels of the target mRNAs of these microRNAs (as specified in TargetScan 7.1; Table S2) in comparison to all other genes. The Cumulative Distribution Function (CDF) plots show that these targets are not changed in proliferation condition, when the three microRNA are not elevated (Figure 3A), however they are specifically upregulated in the tKO cells during differentiation, when the three microRNAs are normally induced (Figure 3B). When the ratio of target : non-target genes were examined in bins of genes distributed from the most-repressed to most-induced in the tKO, the targets were not enriched in any bin in proliferating myoblasts, but were enriched in the bins with induced genes under differentiating condition (bottom plots in Figure 3A and 3B). Thus, the shift for *miR-206/-1a* targets to the right in the CDF plot is due to the upregulation of the target genes in differentiating cells (Figure 3B). Therefore, in tKO C2C12 cells the microRNA targets are de-repressed, suggesting that an unknown regulatory mechanism has not been brought into effect to substitute for the missing microRNAs.

**Figure 3.**
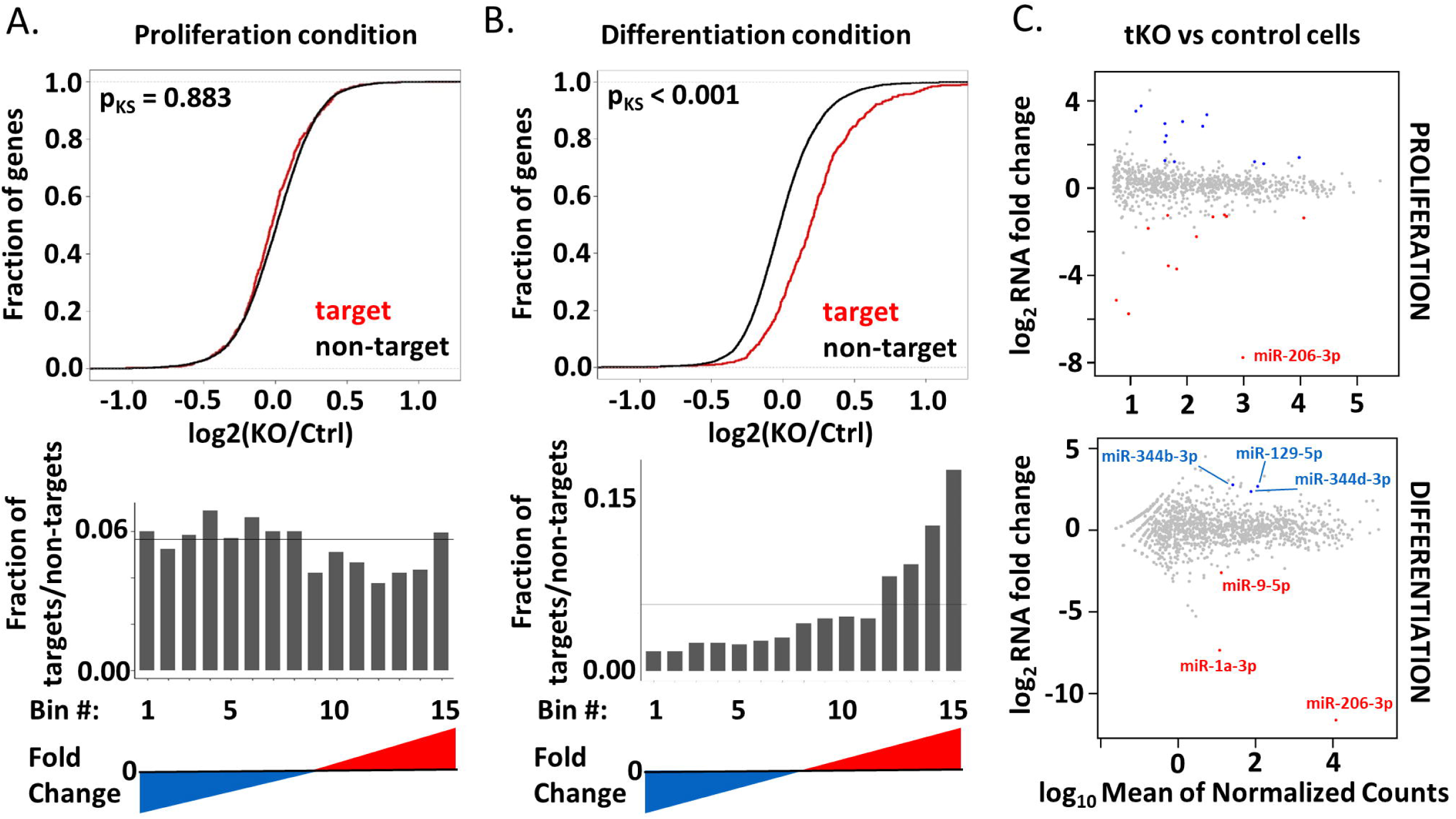
Targets of *miR-206/-1a* are specifically upregulated in differentiating triple KO clones and are not affected by differentially expressed miRNAs. A) Top: Cumulative distribution function (CDF) plot showing no change of the *miR-206/-1* targets in proliferating triple KO clones (KO) versus control cells (Ctrl). The curves for the microRNA targets and non-targets are virtually superimposed. Bottom: Histogram of fraction of the *miR-206/-1* targets, among all bins arranged from the most repressed to most induced in tKO vs Ctrl cells in growth medium. The genes are ranked from the most downregulated (blue) to the most upregulated (red). The horizontal line depicts the uniform distribution expected under the null hypothesis. Targets are based on match to 7mer sequence. B) Top: Cumulative distribution function (CDF) plot showing upregulation of *miR-206/-1* targets in differentiating triple KO clones (KO) versus differentiating control cells (Ctrl). Bottom: Histogram similar to that in Figure 3A, but in differentiating medium. C) MA plots to identify differentially expressed miRNAs between control cells and triple KO clones in proliferation (top) and differentiation conditions (bottom). The microRNA abundance was measured by small RNA-seq. Upregulated microRNAs are presented in blue and downregulated in red (padj<0.1).

To address the hypothesis that other miRNAs could be upregulated in triple KO cells to compensate for the lack of *miR-206/-1a* we performed small RNA-seq in the tKO C2C12 cells. We detect 13 up- and 11 downregulated miRNAs in proliferating tKO cells in comparison to control cells (Figure 3C, S3A, B), and 3 up- and 1 downregulated miRNAs in differentiated tKO in comparison to control cells (Figure 3C, S3C). Nevertheless, the differentially expressed (particularly induced) miRNAs in tKO differentiating cells do not share seed sequence homology with *miR-206/-1a*, and as shown in Figure 3B, do not repress the *miR-206/-1a* targets. Thus it is unlikely that changes in other microRNAs in the tKO cells repress *miR-206/1* targets to compensate for the lack of the latter. We also found the variable change of the linked miR-133 gene product (2-4 fold increase or decrease) which does not meet the FDR < 0.1 threshold (Table S3).

### Lack of miR-206, miR-1a-1 and miR-1a-2 leads to partial embryonic lethality

In order to generate triple KO animals, we first bred single KO of *miR-206*, *miR-1a-1* or *miR-1a-2*, and then their double heterozygous offspring to obtain double KO animals (Figure 4A). Next, we crossed double KO of *miR-206* and *miR-1a-1* animals with double KO of *miR-206* and *miR-1a-2* animals, and subsequently their offspring (*miR-206* KO *miR-1a-1* HET *miR-1a-2* HET) (Figure 4A). The genotypes were confirmed by PCR. Out of 127 genotyped animals we obtained 2 triple KO males. The expected probability of tKO mice was 6.25% and the observed one was 1.57% (Figure 4B), which means the lack of these three miRNAs leads to partial embryonic lethality. This partial embryonic lethality is probably due to the absence of *miR-1a* in the heart, based on previously reported lethality of *miR-1a-1 miR-1a-2* double KO animals (Heidersbach et al., 2013; Wystub et al., 2013) and the embryonic lethality we observe for *miR-1a-1* and *miR-1a-2* double KO animals. Lack of *miR-206* and *miR-1a* expression in the skeletal muscle of the tKO animals was confirmed by q-RT-PCR (Figure 4C).

**Figure 4.**
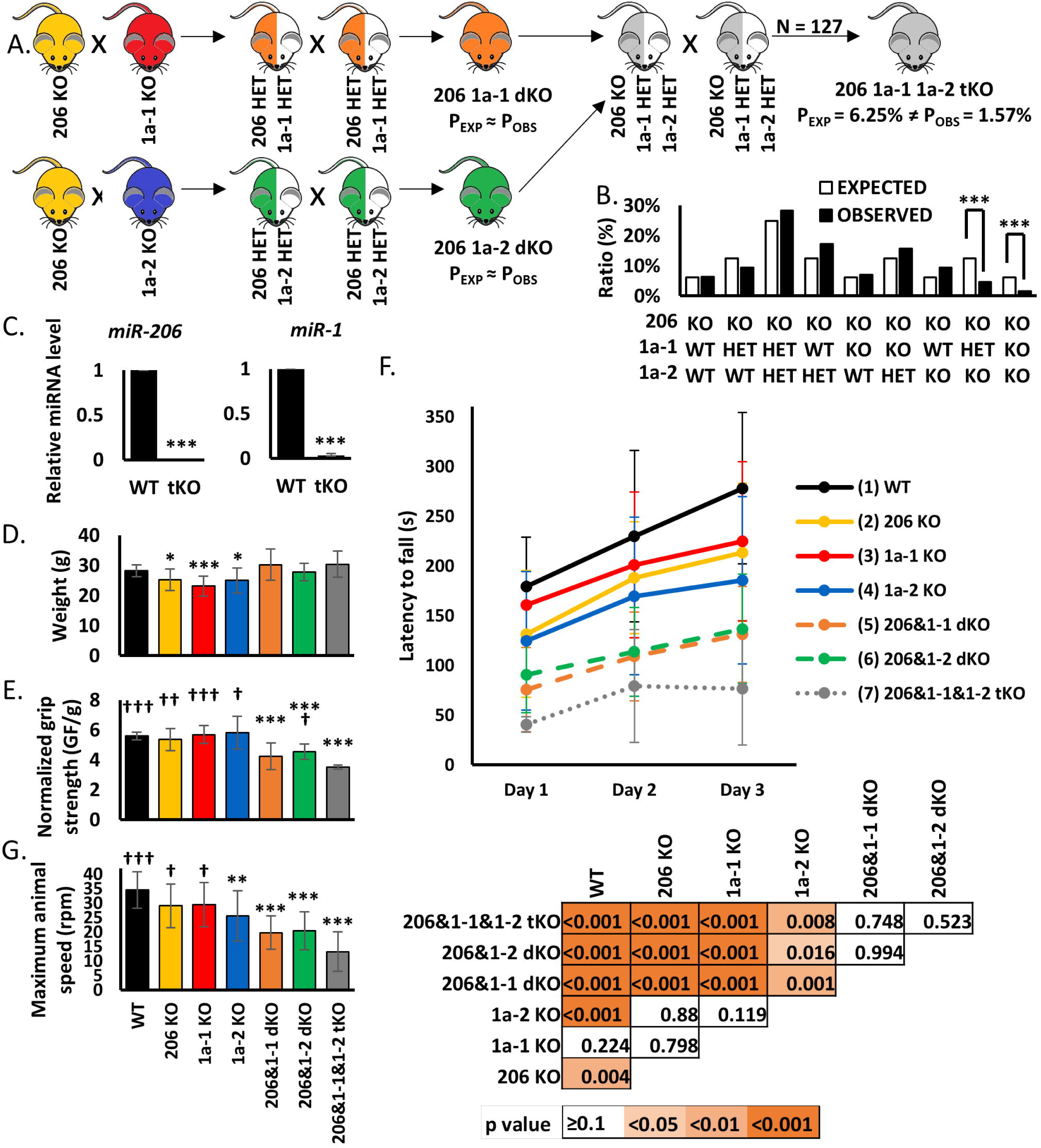
Knockout of *miR-206*, *miR-1a-1* and *miR-1a-2* leads to partial embryonic lethality and diminishes adult mice physical potential. A) Scheme of animal crosses leading to generation of triple KO mice. B) Genotypes of offspring generated from *miR-206* KO *miR-1a-1* HET *miR-1a-2* intercrosses. In total 127 animals were genotyped. Percentage of expected and observed genotypes are given for weaning-age (3-week-old) pups. (***) indicates p-value =< 0.001 C) qRT-PCR analysis of microRNAs in TA skeletal; muscles from control wild-type (WT, n=3) and triple KO (tKO, n=2) animals. Levels of miRNAs were normalized to *U6* snRNA. Levels are shown relative to WT. Values presented as mean +/− SD. Statistical significance was calculated using t-student test. (***) indicates p-value =< 0.001. D) Animal weight used for physical performance tests. Statistical significance was calculated using t-student test. (***) indicates p-value =< 0.001. (*) indicates p-value =< 0.05. N = 13 for all genotypes, except for *miR-206&1-1&1-2* tKO (N = 2). E) Forelimb grip strength measured using grid normalized to respective body weights. Values presented as mean +/− SD. Statistical significance was calculated using t-student test. (***) indicates p-value =< 0.001 in comparison to WT mice. (†††), (††), (†) indicates p-value =< 0.001, =<0.01, =< 0.05 respectively in comparison to tKO mice. N = 13 for all genotypes, except for *miR-206&1-1&1-2* tKO (N = 2). F) Top: Latency to fall measured using rotating rod. Values presented as mean +/− SD. N = 13 for all genotypes, except for *miR-206&1-1&1-2* tKO (N = 2). Bottom: Statistical significance heatmap calculated using one-way ANOVA test. G) Maximum speed of rotation tolerated by the animals measured using rotating rod at the end of experiment (3^rd^ day). Values presented as mean +/− SD. Statistical significance was calculated using t-student test. (***), (**) indicates p-value =< 0.001, =<0.01 respectively in comparison to WT mice. (†††), (†) indicates p-value =< 0.001, =< 0.05 respectively in comparison to tKO mice. N = 13 for all genotypes, except for *miR-206&1-1&1-2* tKO (N = 2).

### Adult triple KO animals have worse physical performance than control mice

Adult triple KO (tKO) mice had the same body weight as the control mice (Figure 4D). To test skeletal muscle function of the tKO animals we measured their grip strength in comparison to control, single KO and double KO mice. Even though the weights of the animals are very similar, there are differences in the force generated. The triple KOs are the weakest animals and this difference is significant in comparison to control, all single KOs and *miR-206 miR-1a-2* double KO animals (Figure 4E). *miR-206 miR-1a-1* double KO and *miR-206 miR-1a-2* double KO were also significantly weaker than control animals (Figure 4E). The Rotarod experiments reveals that the tKO animals fall off the rotarod at an earlier acceleration than the other animal groups. They not only perform significantly worse in this test than control and all single KO animals, but also do not improve between Day 2 and Day 3 as do the other animals (Figure 4F). The tKO mice also have the lowest maximal speed on the third day of experiment of all animals tested (Figure 4G). The progressive decrease in performance from the single KO animals to the double KO animals and further decrease in the triple KO animals confirms that the three microRNAs do compensate for each other and are functionally redundant in skeletal muscle.

### Skeletal muscles and hearts of triple KO animals reveal morphological abnormalities

Even though the quadriceps size is very similar in all experimental mice (Figure 5A), the H&E staining shows that the average fiber cross-sectional area in the tKO animals lacking *miR-206* and *miR-1a* is significantly smaller than in wild type mice (Figure 5B, 5C). Based on previous literature report (Wüst et al., 2018) we decided to check the mitochondria content and organization in muscle fibers. Staining with anti-mitochondria antibody (Figure 5D) does not reveal any differences in fiber type content (Figure 5E). We also did not see a change in number of nuclei visible per fiber cross-section (Figure 5F). Moreover, we do not observe mitochondrial aggregates in the center of fibers or granulate-like patterns in tKO skeletal muscle fibers, phenotypes that were seen in *miR-1a/-133a* knockouts (Wüst et al., 2018). The size of heart in the triple KO animals is comparable to that in wild type (Figure S4A). Masson’s trichrome staining shows accumulation of fibrotic tissue in tKO, but not *miR-206 miR-1a-1* dKO or *miR-206 miR-1a-2* dKO hearts (Figure S4B).

**Figure 5.**
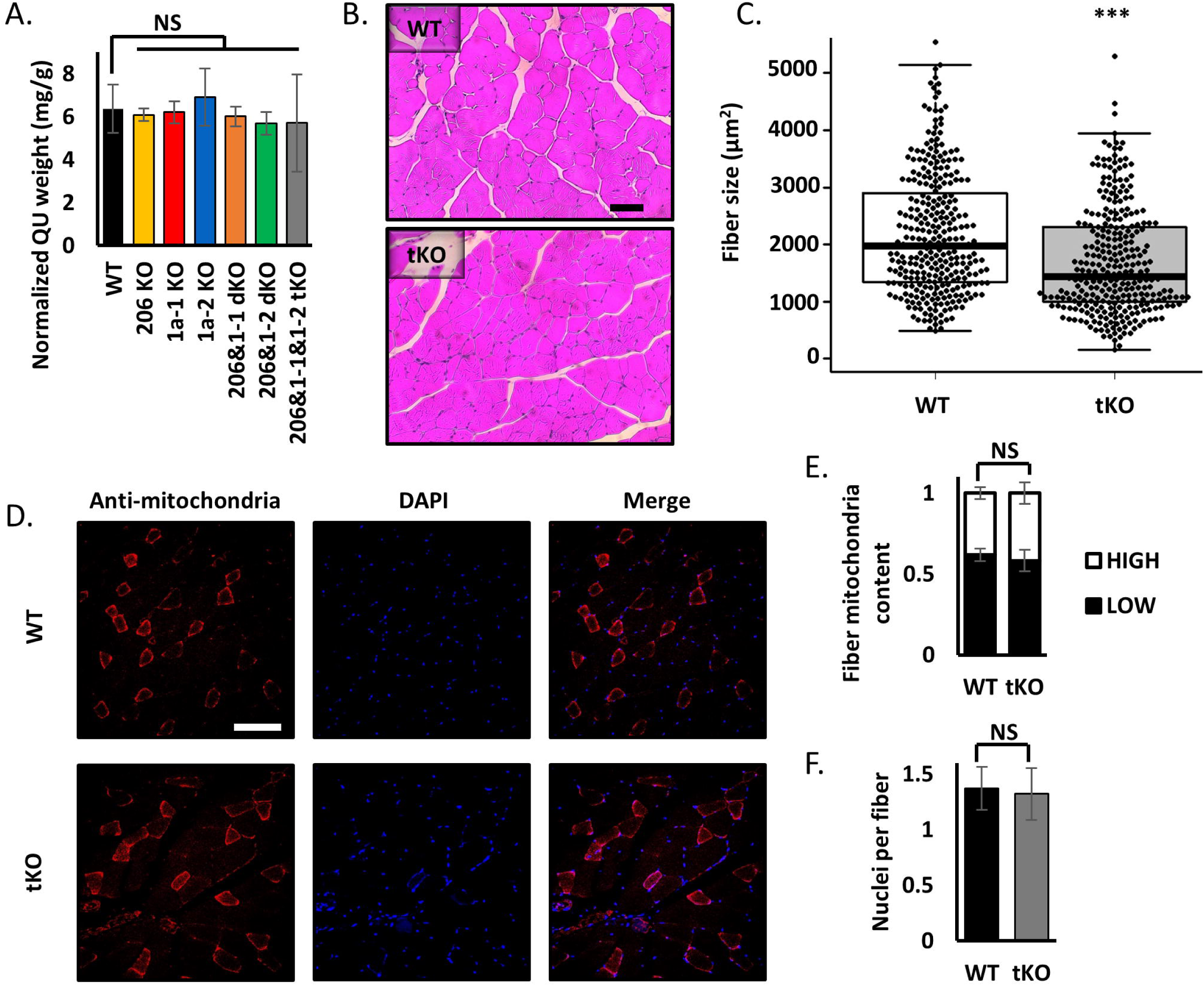
Knockout of *miR-206*, *miR-1a-1* and *miR-1a-2* leads to smaller muscle fiber diameter, but does not alter the content of mitochondria or nuclei in skeletal muscles. A) Quadriceps muscle (QU) weight from animals used in this study. Statistical significance was calculated using t-student test. N = 5 for all group, except *miR-206&1-1&1-2 tKO*, where N=2. B) Representative picture of hematoxylin and eosin (H&E) staining of quadriceps muscle cross-section from WT and tKO mice. Black line = 100µm. C) Quantification of average fiber size based on H&E staining. Statistical significance was calculated using t-student test. (***) indicates p-value =< 0.001. N = 300 fibers per group. D) Representative picture of anti-mitochondria staining of quadriceps muscle cross-section from WT and tKO mice. White line = 200µm. E) Quantification of fiber mitochondria content based on anti-mitochondria staining. High mitochondria content fibers represent type I and IIa, low – type IIb and IId. Statistical significance was calculated using t-student test. N = 600 fibers per group. F) Quantification of nuclei per fiber number based on DAPI (nuclei) and anti-mitochondria (fibers) staining. Statistical significance was calculated using t-student test. N = 600 fibers per group.

## Discussion

Over the last decade many studies showed that myomiRs are important for muscle development and function (Callis et al., 2007; McCarthy, 2011; Townley-Tilson et al., 2010). *miR-206*, *miR-1a-1* and *miR-1a-2* are genes encoding mature *miR-206* and *miR-1a*, muscle specific miRNAs (myomiRs), which not only have identical seed sequence, suggesting functional redundancy, but also are highly expressed in cardiac and skeletal muscle along with *miR-133a* and *miR-133b*. *miR206*, *miR-1a-1* and *miR-1a-2* were suggested to be a critical factor for skeletal muscle differentiation (Anderson et al., 2006; Chen et al., 2006; Chen et al., 2010; Dey et al., 2011; Gagan et al., 2012; Goljanek-Whysall et al., 2012; Heidersbach et al., 2013; Hirai et al., 2010; Kim et al., 2006; Koutsoulidou et al., 2011; Kwon et al., 2005; Mishima et al., 2009; Rao et al., 2006; Sokol and Ambros, 2005; Sweetman et al., 2008; Vergara et al., 2018; Wystub et al., 2013; Wüst et al., 2018; Yuasa et al., 2008; Zhao et al., 2007; Zhao et al., 2005). However, there was no model where all three microRNA genes are knocked out to prove the essentiality of these myomiRs in myogenesis.

Our study argues against an essential role of *miR-206*, *miR-1a-1* and *miR-1a-2* in skeletal muscle differentiation, but supports their functional importance in optimal skeletal muscle differentiation and performance. C2C12 myoblasts lacking *miR-206*, *miR-1a-1* and *miR-1a-2* are still able to differentiate, producing promyogenic mRNAs and proteins, and forming normal myotubes. These findings are true both for PAX7 negative and PAX7 positive murine myoblasts. Although differentiation was observed, in our RNA-seq there was a decrease in expression of genes in mitochondria- and skeletal muscle - related pathways in the triple KO C2C12 clones’ during differentiation. RNA-seq based screens revealed that the targets of these microRNAs remain de-repressed. Thus the simultaneous knockout of the three microRNAs has the expected de-repression of target genes and yet they are not essential for myogenesis. Loss of *miR-1a/-133a* leads to induction of *Mef2a* expression and further induction of *Dlk1-Dio3* mega gene cluster(Wüst et al., 2018). In our *miR-206 miR-1a-1 miR-1a-2* triple KO model despite the decrease of *miR-1a*, *Mef2a* level is not changed in comparison to control cells, and the *Dlk1-Dio3* gene cluster miRNA are not induced. This suggests that the changes seen in Wüst et al. (2018) must be due to the loss of *miR-133a*.

*miR-1* was previously reported as a mitochondrial translation enhancer during muscle differentiation (Zhang et al., 2014). It enters the mitochondria and together with Ago2 but not GW182 stimulates the translation of specific mitochondrial genome-encoded transcripts. The triple KO C2C12 cells show a decrease in the maximal oxygen consumption rate suggesting mitochondrial dysfunction. The mitochondrial respiratory capacity in tKO C2C12 clones is significantly lower than in control C2C12 myoblasts and myotubes. Interestingly, Wüst et al. (Wüst et al., 2018) observed mitochondrial aggregation and accumulation in the skeletal muscle fibers of *miR-1a/-133a* Pax7^+^-Cre mice, whereas we do not see changes in mitochondria structure or localization in our triple KO cells/muscles, suggesting that this phenotype may be caused by lack of *miR-133a*. Therefore, the *miR-206* family (in this report) and *miR-133a* (Wüst et al., 2018) are independently required for normal mitochondrial function.

An *in vivo* model was generated by breeding single knockout animals in order to obtain animals lacking two or three miRNA genes. We obtained *miR-206 miR-1a-1* and *miR-206 miR-1a-2* double knockout animals with ratios similar to expected Mendelian ratios. *miR-1a-1 miR-1a-2* dKO animals were embryonically lethal, most likely due to an essential function of these microRNAs in cardiac differentiation, as published previously (Heidersbach et al., 2013; Wystub et al., 2013). Unexpectedly, we successfully generated triple knockout animals, clearly showing that these miRNAs are collectively not essential for skeletal muscle formation. We still observe the partial embryonic lethality in the triple KO, but we hypothesize that tKO could be generated in a mixed genetic background, which may decrease embryonic lethality, as reported for miR-1a-1 single KO animals by Heidersbach (2013). We hypothesize the triple KO males were infertile as even though breeding was observed, none of the females got pregnant. Interestingly, the mice lacking *miR-206, miR-1a-1 and miR-1a-2* had worse physical performance than wild type, single miRNA knockout and double miRNAs knockout animals. The triple KO mice have weaker muscles and perform worse at rotarod test than wild type control animals. Although the rotarod results may be affected by cardiac defects of KO animals, the decrease in grip strength and decrease in skeletal myofiber diameter clearly show that skeletal muscle function is impaired. Taken together, these findings emphasize that even though *miR-206, miR-1a-1 and miR-1a-2* are not essential for the skeletal muscle formation, they are required for maintenance of its full physical capability. We also found that *miR-206* KO or *miR-1a-2* KO animals perform worse than WT mice in rotarod test, which, to our knowledge, is the first demonstration that even single microRNA gene deletion decreases the neuromuscular performance. This effect increases with each additional microRNA gene deletion.

Extensive cooperation between several miRNAs, mRNAs and proteins leads to effective differentiation. Our study shows that lack of three specific myomiRs *miR-206*, *miR-1a-1* and *miR-1a-2* does not prevent myogenesis, but may lead to functional impairment of skeletal muscles, emphasizing the complex and non-binary nature of skeletal muscle differentiation. Combinatorial knockout of additional known myomiRs may be necessary to fully answer the question if any of these miRNAs are essential for myogenesis. Moreover, this study underscores the importance of joint *in vitro* and *in vivo* full knockout studies by showing that even small alterations in differentiation at the cellular level, may have a significant impact on physical activity with the adult skeletal muscle.

There is an interesting comparison to be made between the functional redundancy of the three myomiRs studied here and of the *MyoD*, *Myf5* and *Mrf4* trascription factor family. Just as overexpression of *miR-206* pushes myoblasts towards differentiation (Kim et al., 2006), overexpression of *Myod1* induces myogenic conversion of fibroblasts (Lattanzi et al., 1998; Tapscott et al., 1988) and ectopic *Myod1* expression drives terminal differentiation of pluripotent stem cells into skeletal myotubes upon chemical treatment (Genovese et al., 2017). *Myod1* knockout *in vitro* inhibits the myoblast differentiation completely (Cichewicz et al., 2018), but *in vivo Myod1* knockout mice develop normal skeletal muscles (Rudnicki et al., 1992). In a similar vein, knockout of individual microRNA genes, *miR-206*, *miR-1a-1* or *miR1a-2*, did not interfere with differentiation in vivo (Heidersbach et al., 2013; Williams et al., 2009; Wystub et al., 2013; Zhao et al., 2007), but affected the regeneration of skeletal muscle after acute or chronic injury (Liu et al., 2012; Williams et al., 2009). However, *Myod1* is dispensable for skeletal muscle development in mice because of the functional redundancy between *Myod1* and *Myf5*, so that mice with double knockout of the two genes do not show skeletal myogenesis (Rudnicki et al., 1993). In a different *Myod1* and *Myf5* double KO model, where *Mrf4* expression was not compromised, the embryonic myogenesis was rescued (Kassar-Duchossoy et al., 2004). Thus the myogenic transcription factors are essential for differentiation, with the essentiality masked by redundancy in actions of the three related transcription factors. This is not the case for the three microRNA genes, where even after removal of all three functionally redundant genes, skeletal muscle differentiation still occurs *in vitro* and *in vivo*: the three microRNAs studied here are not essential for differentiation.

However, the three microRNAs are clearly important for optimal skeletal muscle differentiation, judging by the defects in mitochondrial function, myofiber diameters and skeletal muscle performance that we report in this paper. We also demonstrate that it is possible to genetically separate the functional role of miRNAs coming from bicistronic loci like *miR-1a* and *miR-133a* and to genetically examine functional redundancy of as many as three microRNAs *miR-206*, *miR-1a-1* and *miR-1a-2*.

## Material and Methods

### Cell lines generation

#### Stable overexpression of inducible Cas9 in C2C12 cells

pCW-Cas9 (addgene #50661) vector was packed in the virus using psPAX2 (addgene #12260) and pMD2.G (addgene #12259) in 293T cells. C2C12 cells were transduced with the filtered supernatant containing virus. After 24 hours cells were treated with puromycin (C=2ug/ml; Sigma, Cat# P9620) and resistant cells were seeded to 96-well plates using single cell dilution method. Growing clones were examined for Cas9 expression by immmunoblotting after treatment with doxycycline (C=1ug/ml; Sigma, Cat# D9891) for 48h. Clones with high expression upon induction and low leakage level without doxycycline were next tested for differentiation efficiency by q-RT-PCR for *Myod1*, *Myogenin* and *Myh3*, and immunoblotting for MYOD1, MYOGENIN and MHC. The clone, which was the most similar to wild-type cells, was chosen for further experiments.

#### Knockout cell line generation

CRISPR protocol (Ran et al., 2013) with minor changes was followed to achieve deletion of miR-206, miR-1a-1 and miR-1a-2 genes. Briefly, sgRNAs were designed using CRISPR DESIGN tool: http://crispr.mit.edu/. Cas9 expression in C2C12 cells was induced 24h before sgRNAs transfection using doxycycline (C=1ug/ml). Cells were co-transfected with 6 different sgRNAs cloned into gRNA_GFP-T2 (addgene #41820), and a spiking vector coding for a resistance gene using Lipofectamine 3000 (Life Technologies, Cat# L3000015). After 24-48 hours cells were treated with hygromycin (C=300ug/ml; Life technologies, Cat# 10687-010), and resistant cells were seeded to 96-well plates using single cell dilution method. Growing clones were examined for desired deletion by PCR on extracted genomic DNA (Quick Extract DNA Extraction Solution, Lucigen., Cat# QE09050), and candidates with complete loss of all three WT PCR products (homozygous deletions) were confirmed by Sanger sequencing and q-RT-PCR for *miR-206*, *miR-1a-1* and *miR-1a-2*. Oligonucleotides sequences are listed in Table S1.

### Cell culture, differentiation assay

C2C12 cells were cultured in DMEM-high glucose medium (GE Healthcare Life Sciences co., Cat# SH30022.FS) with 10% FBS (Gibco, Cat# 10437-028), when differentiating, serum was switched to 2% horse serum (GE Healthcare Life Sciences co., Cat# SH30074.03) supplemented with (1) 1x ITS (Insulin-Transferrin-Selenium; Fisher, Cat# 41400045) and 5mM LiCl (Sigma; Cat# 203637) for PAX7 negative cells or (2) 1x ITS for PAX7 positive cells. PAX7 negative and positive cell lines were independently purchased.

### RNA isolation and RT-PCR

RNA was isolated by TRIzol reagent (Life Technologies, Cat# 15596018) using Direct-zol RNA MiniPrep Plus Kit including DNase treatment (Zymo Research, Cat# R2052). cDNA synthesis for mRNA expression levels measurement was performed using Superscript III RT cDNA synthesis kit (Life Technologies co., Cat# 18080-051) with random hexamer and oligodT priming. After cDNA synthesis, qPCR was performed with Applied Biosystems 7500 Real-Time PCR Systems using SensiFAST™ SYBR® Hi-ROX Kit (BIOLINE, Cat# BIO-92020). miScript II RT (Qiagen, Cat# 218161) and miScript SYBR® Green PCR kits (Qiagen, Cat# 218075) were used for miRNA and pre-miRNA expression levels measurement. All primers used in this study are listed in Table S1.

### Western blotting

Cells were lysed in IPH buffer (50mM Tris-Cl, 0.5% NP-40%, 50mM EDTA) and run on 10% polyacrylamide SDS-PAGE gel, transferred to nitrocellulose membranes. Membranes were blocked for 30 minutes in 5% milk containing PBST, and incubated overnight with primary antibody in 1% milk. Secondary antibody incubation was carried out for 1 hour after washing, and at 1:4000 dilution before washing and incubation with Millipore Immobilon HRP substrate. Primary antibodies were used as follows: mouse monoclonal MyoD (5.8A) (Santa Cruz co., Cat# sc-32758), mouse monoclonal Myogenin (F5D) (Santa Cruz co., Cat# sc-12732), rabbit polyclonal MYH3 (Proteintech, Cat# 22287-1-AP), mouse monoclonal HSP90 α/β(F-8) (Santa Cruz co., Cat# sc-13119). Secondary antibodies were used as follows: anti-rabbit IgG, HRP-linked Antibody (Cell Signaling, Cat# 7074S), anti-mouse IgG, HRP-linked Antibody (Cell Signaling, Cat# 7076S).

### Immunofluorescence assay

Cells were plated on glass coverslips and collected in growth medium or after 5 days of differentiation. The coverslips were fixed with 4% paraformaldehyde in PBS for 15 min, permeabilized in 0.5% Triton X-100 in PBS, and blocked in 5% goat serum. The coverslips were incubated with primary rabbit polyclonal antibody MYH3 (Proteintech, Cat# 22287-1-AP) overnight at 4^◦^C and then with goat anti-Rabbit IgG (H+L) Cross-Adsorbed Secondary Antibody, Alexa Fluor 647 (Invitrogen, Cat# A-21244) for 1 h. Cells were stained with Hoechst 33342 (1 µg/mL; Life Technologies, Cat# H3570) for 2 minutes at room temperature, washed and then mounted with ProLong™ Gold Antifade Mountant (Life Technologies, Cat# P10144). The primary and secondary antibodies were diluted 1:400 and 1:1000, respectively. The fusion index was calculated by dividing the number of nuclei contained within multinucleated cells by the number of total nuclei in a field. Microscopy was performed using the Zeiss LSM 710 Confocal Microcopy and ImageJ Software for analysis (Schneider et al., 2012).

### Mitochondrial Stress Test

Oxygen Consumption Rate (OCR) for Mitochondrial Stress Test (MST) assay was measured using Seahorse XF24 Extracellular Flux Analyzer (Agilent). 20,000 cells per well were plated in growth medium in at least triplicate for each cell line and condition 48h before test. For DM1, 24h post seeding medium was switched to differentiation and MST assay was performed after 24h. One hour before MST assay medium was changed to MST medium (unbuffered, serum-free DMEM pH 7.4; Life Technologies, Cat# 12100046). For the MST assay, oligomycin (inhibits ATP synthase; Sigma, Cat# 75351), BAM15 (protonophore uncoupler, stimulates a maximum rate of mitochondrial respiration; Cayman, Cat# 17811), and Rotenone and Antimycin A (inhibits the transfer of electrons in complex I and III, respectively; Sigma, Cat# R8875, Cat# A8674) were injected to a final concentration of 2µM, 10µM, 1µM and 2µM, respectively over a 100-minute time course. At the end of each experiment, each assay was normalized to total protein count measured from a sister plate that was seeded concurrently with the experimental plate.

### RNA-seq

RNA samples were isolated from proliferating or differentiating WT and tKO cells as described above. RNA-seq was performed by Beijing Novogene Co., Ltd. on poly(A) enriched RNA using the Illumina HiSeq instrument. We obtained >50 million paired-end 150 bp long reads for all conditions. RNA-Seq data was aligned to the reference genome - mouse assembly GRCm38/mm10 using STAR v2.5 (Dobin et al., 2013) and quantified by HTSeq 0.6.1p1 (Python 2.7.5) (Anders et al., 2015). DESeq2 R package (Love et al., 2014) was then applied to determine differentially expressed genes with a significant criterion padj <0.05. Gene Ontology was performed using clusterProfiler (Yu et al., 2012). GSEA (Subramanian et al., 2005) was used for gene set enrichment analysis. The list of *miR-*206*/-1a* targets was downloaded from TargetScan 7.1 (Agarwal et al., 2015). All RNA-Seq libraries data files are available under GEO accession number: GSE133260.

### Short RNA-seq

RNA samples were isolated from proliferating or differentiating WT and tKO cells as described above. 0.5 ug of RNA was used for short RNA-seq library preparation with NEBNext Small RNA library kit (New England Biolabs, Cat# E7300). Shortly, 3’ preadenylated adaptors and then 5’ adaptors were ligated to RNA, followed by reverse transcription and PCR with indexes. Next, 8% TBE-PAGE gel was used for size selection (15-50nt). For sequencing we used Illumina NextSeq 500 sequencer with high-output, 75-cycles single-end mode at Genome Analysis and Technology Core (GATC) of University of Virginia, School of Medicine. Data trimmed by catadapt v1.15 (Python 2.7.5) (Martin, 2011) was aligned to the reference genome (gencode GRCm38.p5 Release M16, primary assembly) using STAR v2.5 with settings based on previous paper (Faridani et al., 2016) and the total number of mapped reads were used for normalization. In general, mapped percentage is more than 95%. Unitas v.1.5.1 (with SeqMap v1.0.13) (Gebert et al., 2017; Jiang and Wong, 2008) was used for miRNA abundance quantification with setting –species_miR_only –species mus_musculus to map the reads to mouse sequence of miRBase Release 22 (Kozomara et al., 2019). This setting (equivalent to –tail 2 –intmod 1 –mismatch 1 –insdel 0) will allow 2 non-templated 3’ nucleotides and 1 internal mismatch for miRNA mapping. For differential analysis, DESeq2 (Love et al., 2014) was used on count matrix of mature microRNAs. All short RNA-Seq libraries data files are available under GEO accession number: GSE133255.

### Mice

Mouse strains used in the study were obtained as follows: *miR-206* KO mice provided by Eric Olson (Williams et al., 2009); *miR-1a-1* and *miR-1a-2* KO mice provided by Deepak Srivastava (Heidersbach et al., 2013; Zhao et al., 2007). Double KO mice were generated by crossing single KO animals and then crossing their double heterozygous offspring. Triple KO mice were generated by crossing *miR-206 miR-1a-1* double KO animals with *miR-206 miR-1a-2* double KO animals and then crossing their offspring (*miR-206* KO *miR-1a-1* HET *miR-1a-2* HET) as shown in Figure 4A. All animals were PCR-genotyped using gene-specific primers listed in Table S1. Work involving mice adhered to the guidelines of the University of Virginia Institutional Animal Care and Use Committee (IACUC), protocol number 3774.

### Physical endurance tests

Tests described below were performed on 12 weeks old mice, males and females (7+6) for all animals except tKO mice, where 2 males were examined.

#### Grip strength test

Mice were tested for their maximal grip strength with a grip strength meter (C.S.C. Force Measurement, Inc. AMETEK 2LBF Chatillon). The mice were placed on the force meter allowing forelimb to grip the grid. The mice were then slowly dragged backward until loss of grip. The bodies of the mice were kept in a horizontal position during the test. Three trials were repeated in 1-minute intervals. The average of the three trials was recorded as the maximum grip strength.

#### Accelerating rotarod test

Motor coordination and balance were assessed on an Economex accelerating rotarod (Columbus Instruments) that has the capacity to test five mice simultaneously. The testing procedure consists of three days, two runs on each day. The rod was adjusted to spin at a beginning speed of 4.0 r.p.m with an acceleration (0.12 r.p.m.) over 300s to final speed 40.0rpm. The maximum running time was 360s. Latency to fall from the rotarod and maximum animal speed for each run were recorded and averaged per each day.

Experiments involving mice adhered to the guidelines of the University of Virginia Institutional Animal Care and Use Committee (IACUC), protocol number 4064.

### Muscle isolation, histological analyses and staining

The mice were sacrificed by carbon dioxide inhalation. The hearts, quadriceps and tibialis anterior muscles were dissected quickly, weighed, fixed with 4% paraformaldehyde or snap-frozen in liquid nitrogen and stored at −80 °C. All samples were collected at least 7 days after assessment of physical function from 3 males and 2 females for all animals except tKO mice (2 males)

All formalin-fixed paraffin-embedded (FFPE) sections, H&E staining and Masson’s trichrome staining were performed by Research Histology Core at University of Virginia (Charlottesville, USA). Fiber size was measured on H&E stained quadriceps muscles using ImageJ 1.50i (Java 1.6.0_24) (Schneider et al., 2012) on three pictures from two animals per genotype (300 fibers per genotype in total).

Immunofluorescence on FFPE sections was performed as previously described (Wang et al., 2014). In short, FFPE sections were deparaffinized and antigen-retrieved (by Biorepository & Tissue Research Facility at University of Virginia), washed, and blocked then incubated with anti-mitochondria antibody, clone 113-1, Cy3 conjugate (EMD Millipore, Cat# MAB1273C3) overnight. Slides were incubated with CuSO4 to reduce autofluorescence and mounted with Prolong Gold antifade reagent with DAPI. Images were taken with a Zeiss LSM 710 Multiphoton microscope with a 20× (NA 0.8) objective. Mitochondria content and nuclei per fiber count was performed using ImageJ 1.50i (Java 1.6.0_24) (Schneider et al., 2012) on ten pictures per genotype (600 fibers per genotype in total).

## Acknowledgments

We thank all members of the Dutta lab for helpful discussion, especially Dr. Etsuko Shibata for technical guidance regarding CRISPR/Cas9 method. We thank Research Histology Core, Genome Analysis and Technology Core, and Biorepository & Tissue Research Facility at University of Virginia (Charlottesville, USA). We thank Scott Zeitlin and Elise Braatz for help with grip strength measurement, and Ashley Bolte for help with rotarod experiment. This work was supported by a grant from the NIH (RO1 AR067712) (AD). RKP was supported by a Predoctoral Fellowship from the American Heart Association (18PRE33990261). The authors declare no competing financial interests.

## Author Contributions

RKP and AD designed all experiments. RKP obtained all C2C12 cell lines and mice strains used in this study. RKP and ES confirmed deletions in KO clones. RKP and ES performed differentiation assays, qPCR and Western Blot analyses for C2C12 cells. ES performed immunofluorescence experiments. RKP prepared samples for RNA-seq and short RNA-seq experiments. ZS performed and analyzed short RNA-seq experiment. RKP crossed and genotyped all animals with help of KJ. Physical endurance experiments were performed by RKP. Muscle isolation was done by RKP with help of PP and KJ. SN performed Seahorse assay with help of RKP. JK performed anti-mitochondria staining. JL helped with mice study design and DK with mitochondria-related study design. RKP and AD wrote the manuscript.

## Supplemental item titles

*Table S1.* Oligonucleotides used in this study.

*Table S2.* Expression of *miR-206/1a* targets based on RNA-seq results.

*Table S3.* Expression of *miR-133* family based on small RNA-seq results.

## Supplemental figure legend

**Supplemental figure 1.**
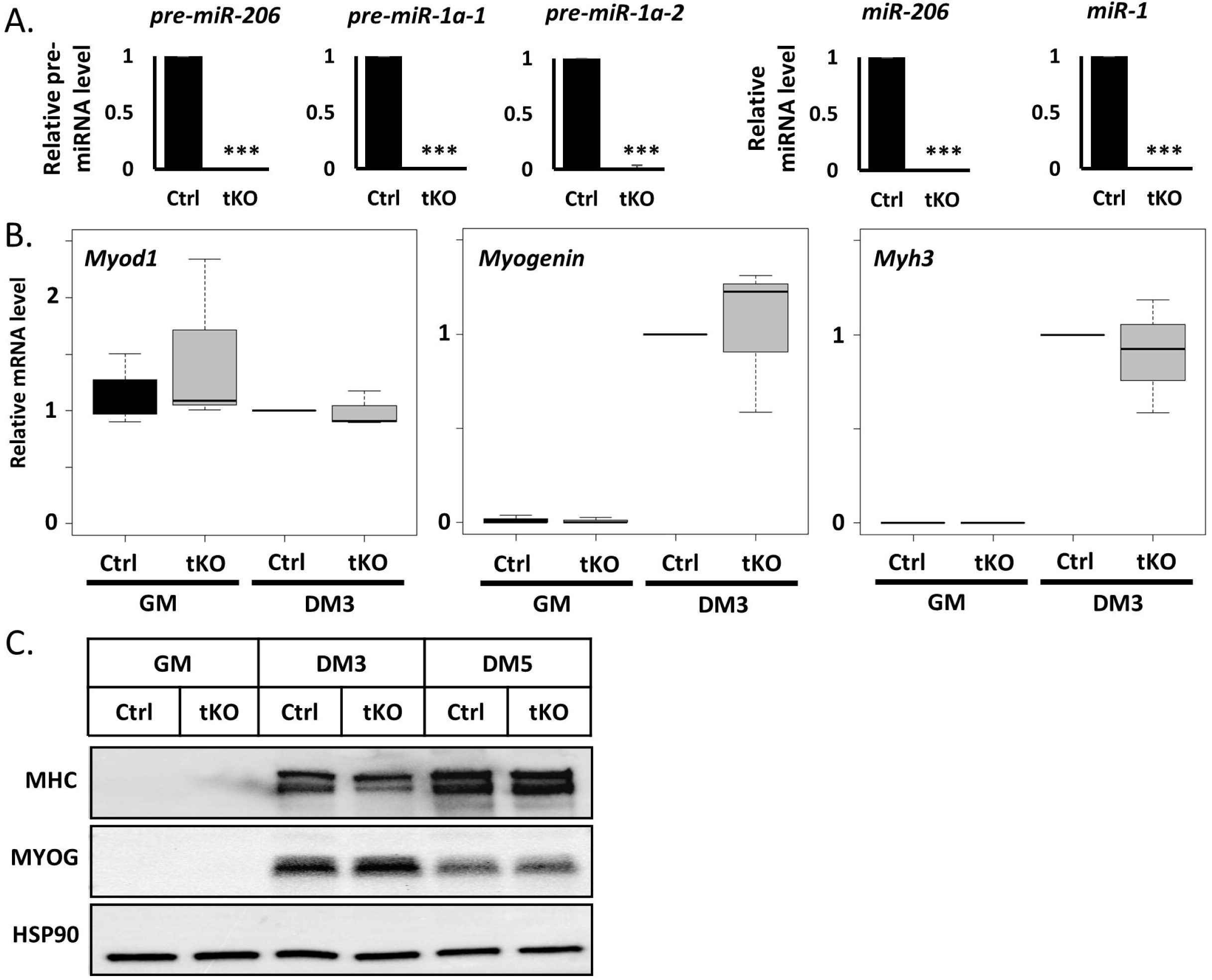
Simultaneous knockout of miR-206, miR-1a-1 and miR-1a-2 genes in PAX7 positive murine myoblast doesn’t block differentiation, but slightly decreases MHC protein level. A) qRT-PCR analysis of differentiating (DM3) control cells (Ctrl) and triple KO clones (tKO C1 and tKO C2). Levels of pre-miRNAs were normalized to Gapdh and miRNAs – to U6 snRNA. Levels are shown relative to control cells (Ctrl DM3). Values represent three biological replicates, presented as mean +/− SEM. Statistical significance was calculated using t-student test. (***) indicates p-value =< 0.001. B) qRT-PCR analysis of proliferating (GM) and differentiating (DM3) control cells (Ctrl) and triple KO clones (tKO C1 and tKO C2). Levels of Myod1, Myogenin and Myh3 mRNAs were normalized to Gapdh and shown relative to control cells (Ctrl DM3). Box and whiskers plot from three biological replicates with black line representing the median. Statistical significance was calculated using t-student test. C) Western blot of proliferating (GM) and differentiating (DM1, DM3, DM5) control cells (Ctrl) and triple KO clones (tKO C1 and tKO C2). Protein levels for MYOGENIN and MHC were measured. HSP90 serves as a loading control.

**Supplemental figure 2.**
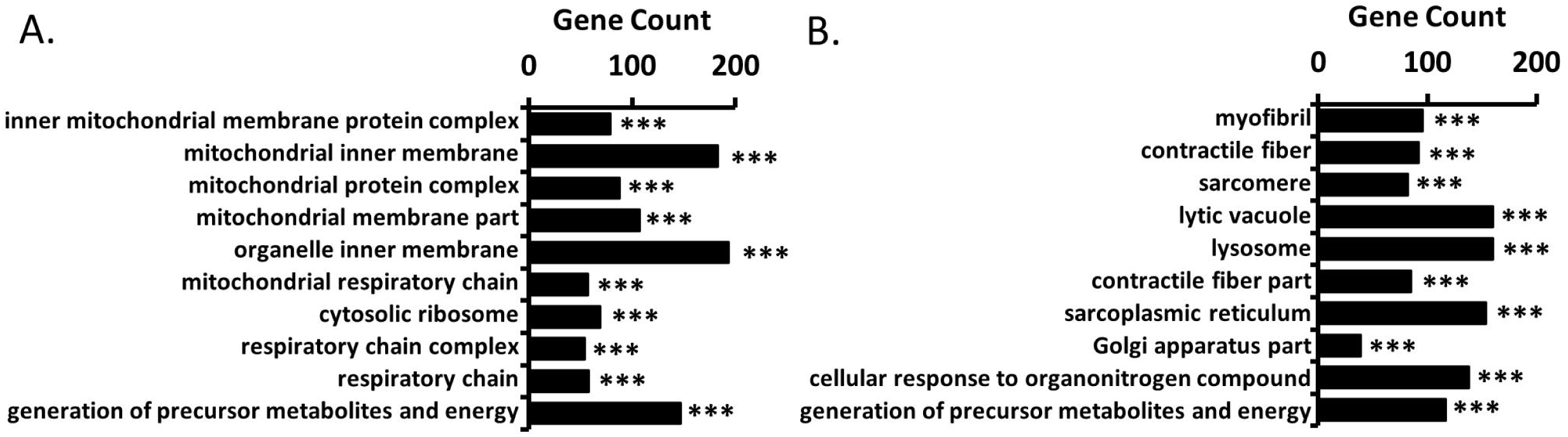
Lack of miR-206/-1a leads to mitochondria function impairment in C2C12 cells. A) Top 10 significant Gene Ontology terms enriched in genes upregulated during differentiation of control cells (Ctrl). (***) indicates padj =< 0.001 B) Top 10 significant Gene Ontology terms enriched in genes upregulated during differentiation of triple KO clones (tKO1 and tKO2). (***) indicates padj =< 0.001

**Supplemental figure 3.**
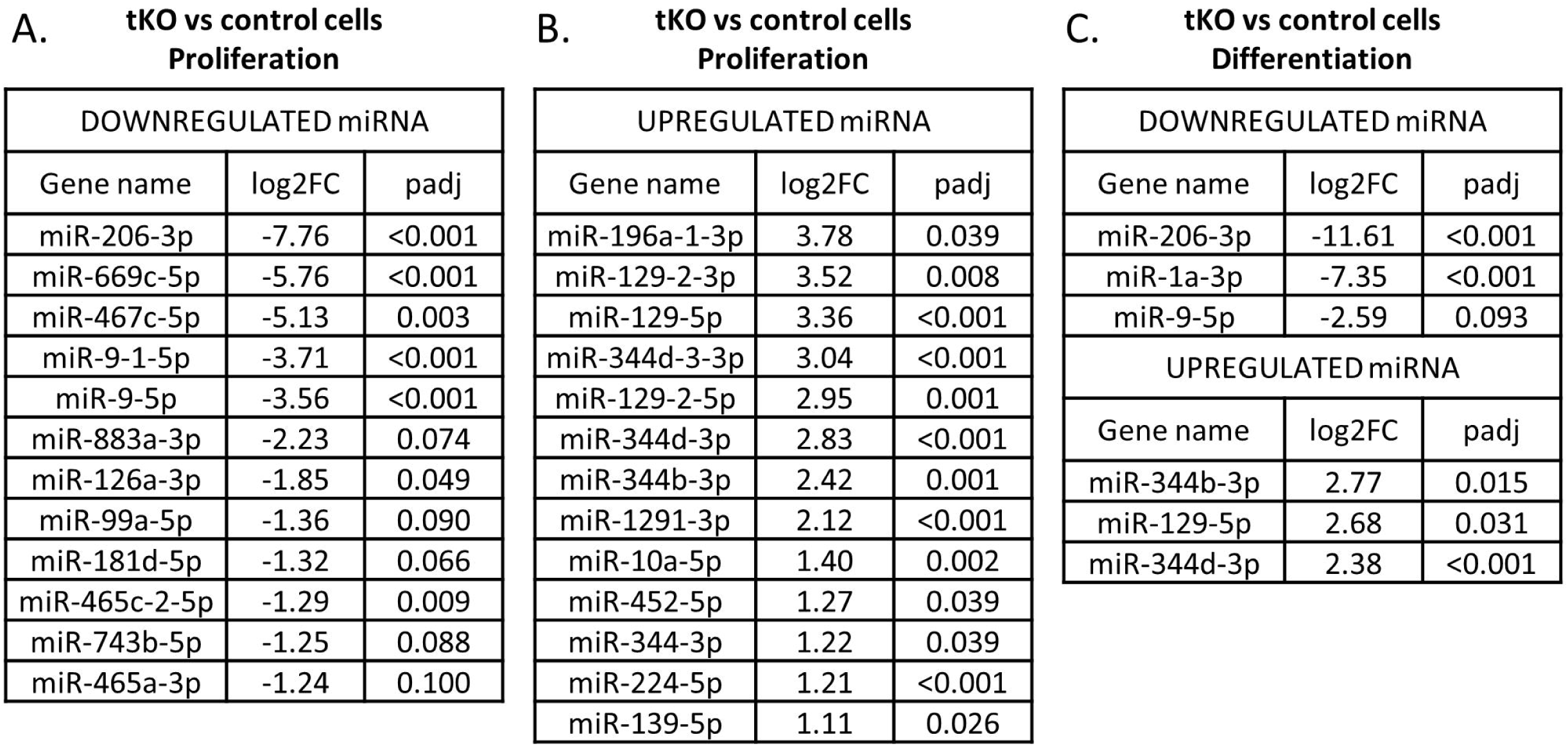
Differentially expressed miRNAs in triple KO cells do not share seed-sequence with miR-206/-1a and so are unlikely to repress miR-206/-1a targets. A) miRNAs downregulated in proliferating triple KO clones versus control cells. padj =< 0.1 B) miRNAs upregulated in proliferating triple KO clones versus control cells. padj =< 0.1 C) miRNAs down- and upregulated in differentiating triple KO clones versus control cells. padj =< 0.1

**Supplemental figure 4.**
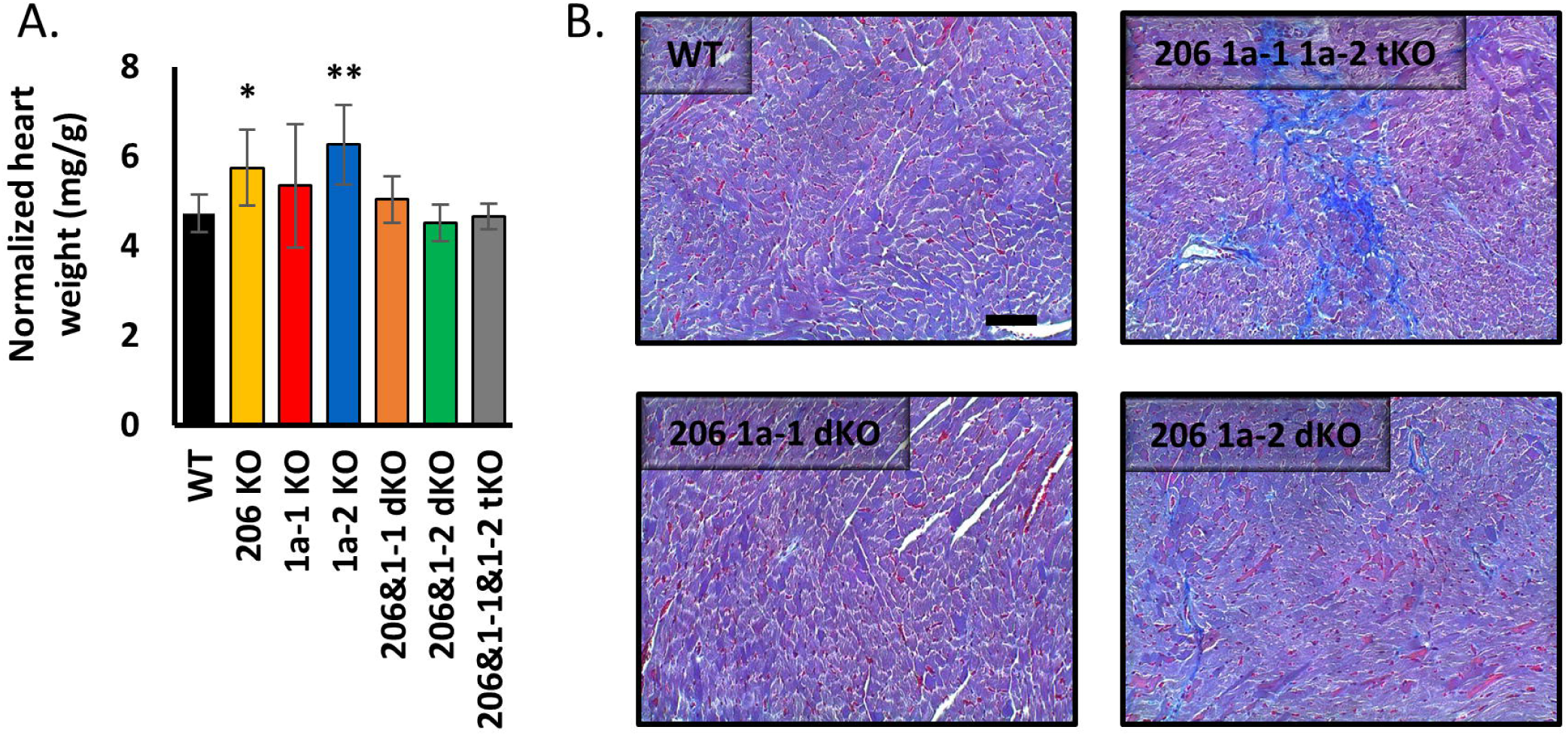
Mice with triple knockout of miR-206, miR-1a-1 and miR-1a-2 show high levels of heart fibrosis. A) Hearts weight from animals used in this study. Statistical significance was calculated using t-student test. N = 5 for all group, except miR-206&1-1&1-2 tKO, where N=2. B) Representative picture of Masson’s trichrome staining of heart section from WT, miR-206 miR-1a-1 dKO, miR-206 miR-1a-2 dKO and miR-206 miR-1a-1 miR-1a-2 tKO mice. Black line = 100µm

